# Development and application of species ID and insecticide resistance assays, for monitoring sand fly *Leishmania* vectors in the Mediterranean basin and in the Middle East

**DOI:** 10.1101/2024.07.30.605769

**Authors:** Sofia Balaska, Jahangir Khajehali, Konstantinos Mavridis, Mustafa Akiner, Kyriaki Maria Papapostolou, Latifa Remadi, Ilias Kioulos, Michail Miaoulis, Emmanouil Alexandros Fotakis, Alexandra Chaskopoulou, John Vontas

## Abstract

**Background:** Development of insecticide resistance (IR) in sand fly populations is an upcoming issue of public health concern, threatening leishmaniasis mitigation efforts by insecticide- based vector control. There is a major knowledge gap in the IR status of wild populations worldwide, possibly attributed to the unavailability of specialized tools, such as bioassay protocols, species baseline susceptibility to insecticides and molecular markers, to monitor such phenomena in sand flies.

**Methodology/ Principal findings:** Several sand fly populations from (semi-)rural regions of Greece, Turkey and Iran were sampled and identified to species, showing populations’ structure in accordance with previously reported data. Genotyping of known pyrethroid resistance-associated loci revealed the occurrence of voltage-gated sodium channel (*vgsc*) mutations in all surveyed countries. Knock-down resistance (*kdr*) mutation L1014F was prevalent in Turkish regions and L1014F and L1014S were recorded for the first time in Iran and in Turkey and Greece, respectively, yet in low frequencies. Moreover, CDC bottle bioassays against pyrethroids in mixed species populations from Greece indicated full susceptibility, using though the mosquito discriminating doses. In parallel, we established a novel individual bioassay protocol and applied it comparatively among distinct *Phlebotomus* species’ populations, to detect any possible divergent species-specific response to insecticides. Indeed, a significantly different knock-down rate between *P. simici* and *P. perfiliewi* was observed upon exposure to deltamethrin.

**Conclusions/Significance:** IR in sand flies is increasingly reported in leishmaniasis endemic regions, highlighting the necessity to generate additional monitoring tools, that could be implemented in relevant eco-epidemiological settings, in the context of IR management. Our molecular and phenotypic data add to the IR map in a macroarea with otherwise limited data coverage.

**Author Summary:** Phlebotomine sand flies vector leishmaniasis to human and animals, a neglected tropical disease of the (sub-)tropics, displaying an expanding epidemiological tendency towards previously non-endemic regions. As leishmaniasis eradication largely relies on vectors’ insecticidal control, regular monitoring of insecticide resistance (IR) is a core element of integrated vector management. IR data are limited in sand fly populations worldwide though, among others due to the unavailability of robust molecular diagnostic and phenotyping tools. Here, we aim to characterise the pyrethroid resistance (PyR) status of populations originating from countries of the Mediterranean basin and the Middle East; i.e. Greece, Turkey and Iran. PyR-associated mutations were detected in all three countries, while the populations from Greece exhibited susceptibility upon exposure to deltamethrin in CDC bioassays. We established an individual bioassay protocol to enable IR monitoring in settings with multi-species population structure, and applied it comparatively among three distinct *Phlebotomus* species, revealing that they respond differently to insecticide treatment. Our study will facilitate systematic surveillance and evidence-based sand fly control in the endemic regions.

## Introduction

Leishmaniasis is classified among the top ten epidemiologically impacting neglected tropical diseases globally, causing approximately 30,000 parasite-causative fatalities out of almost one million cases annually [1]. Phlebotomine sand flies mediate the transmission of *Leishmania* parasites to humans and animals, in the tropical and sub-tropical zones around the world.

Even though low- and middle-income countries are disproportionately burdened by leishmaniases occurrence [1], the epidemiological scenarios in central and southern Europe and the Mediterranean basin have been changing, over the last decades, with new disease foci appearing in previously non-endemic regions [2–4]. Climatic crisis, facilitating the spread of sand flies in cooler environments, concurrent with environmental modifications/urbanization and population movement (i.e. migration/ mass population displacement, tourism, travelling usually along with reservoir hosts, as for example dogs) [5], possibly trigger this epidemiological tendency [6].

The role of vector control to combat leishmaniasis transmission is crucial, especially due to the unavailability of vaccines for humans, as well as several issues accompanying antileishmanial chemotherapy (e.g. drug resistance, toxicity, high cost) [7]. Insecticide-based methods (i.e. indoor residual spraying, insecticide-treated bed nets, insecticide durable wall lining, etc) have achieved focal reduction of sand fly populations in highly endemic regions, yet primarily within the framework of mosquito vector control programs, rather than targeted regional sand fly control campaigns [8]. Nevertheless, the selection pressure imposed by intensive insecticide applications for public health protection and/or off-target effects from agriculture results in the development of insecticide resistance (IR) in sand flies [9, 10]. There is fragmented assessment of sand fly vectors resistance to insecticides and limited molecular data for the underlying resistance mechanisms. The majority of those resistance records are aggregated in leishmaniasis hyper-endemic regions of southeastern Asia and the Middle East, and refer mainly to DDT and pyrethroids [11], with the latter being the primary class of insecticides deployed in public health interventions [10]. Resistance phenotypes have been often associated with the presence of the voltage-gated sodium channel (*vgsc*) mutations L1014F/S (knock-down resistance mutations; *kdr*) [12, 13].

At present, there are limited IR monitoring tools specialized for sand flies. Only recently, WHO established the discriminating concentrations for WHO tube bioassays against currently used insecticides in five phlebotomine species (i.e. *P. papatasi, P. longipes, P. duboscqi, P. argentipes* and *L. longipalpis*) [14, 15], while no specific CDC bottle bioassay guidelines are available [16]. A few studies so far have been conducted to assess the baseline susceptibility levels in CDC bioassays, often with contradictory results [17–19]; nevertheless, fundamental data for most sand fly vector species are missing. Furthermore, current bioassays for monitoring phenotypic resistance in wild vector populations (WHO tube, CDC bottle) are based on pooling samples of same vector species in single vials/tubes [14, 15, 20]. While for mosquitoes this is plausible since species discrimination relies on external morphology and wild-caught specimens can quickly and easily be grouped to genus and/or species, it may not readily apply to sand flies. The reason being that sand fly identification depends on internal morphology requiring dissection and mounting of specimens, thus current IR monitoring protocols need to be adjusted/adapted to allow for discriminating sand fly species when collections are made in regions with mixed sand fly species composition.

Contrary to mosquito vectors, where research has revealed a variety of IR mechanisms (i.e. target-site mutations, elevated detoxification, reduced cuticle penetration, behavioural avoidance and insecticide sequestration or excretion) [21–24], the paucity of genomic resources for sand flies seriously impedes deep investigation of such molecular basis, as well as the development of relevant diagnostic markers. All the above limitations critically deter regular IR monitoring, hence, evidence-based management efforts in sand fly populations.

In order to enhance the scientific understanding of IR in sand fly populations in areas with limited data, we focus on the area of the eastern Mediterranean basin and the Middle East, particularly on Greece, Turkey and Iran. All three countries share a long record of autochthonous leishmaniases transmission, while last decade’s epidemiological data denote an escalating trend of visceral (VL) and cutaneous leishmaniasis (CL) prevalence, in Greece and in Turkey and Iran, respectively [25–27]. Despite the rich sand fly diversity, certain vector species are more widely distributed and primarily implicated in parasite circulation in each country. *Phlebotomus perfiliewi, P. tobbi* and *P. neglectus* mediate *L. infantum* transmission in Greece [28], while *P. alexandri* is also dominant along with those aforementioned VL vectors in Turkey [29]. *Phlebotomus papatasi* and *P. sergenti* serve as the main vectors of CL causative agents, *L. major* and *L. tropica*, in Turkey and Iran [29–31].

Here, our main objectives included: i) monitoring sand fly species composition in populations collected from 12 regions of Greece, Turkey and Iran; ii) analysing pyrethroid resistance associated *Vgsc* gene loci in these populations; iii) phenotypically assessing by CDC bottle bioassays the response to deltamethrin of three mixed species sand fly populations from Greece; iv) designing a novel individual-specimen bioassay (a modified version of the CDC bottle bioassay), proposed to operate in regions with mixed sand fly species composition, and applying it here to compare three distinct *Phlebotomus* species against deltamethrin. Our molecular and bioassay data enrich the scientific evidence for the IR status of sand fly populations in areas with poor data coverage, thereby support evidence-based vector control efforts.

## Materials and Methods

### 1. Sampling areas, sand fly collection and sample handling

Multiple sand fly collections were performed in regions of Greece, Turkey and Iran, between 2019 and 2023. Locations were selected in the interface between semi-rural environment in agricultural areas and urbanized territories (e.g. residencies, streets), occasionally in close proximity to animal farms (Table 1; Figure 1). Samples were collected from at least 2 to 3 different sites in each location, to avoid the probability of including isofemale sand flies in the downstream analyses. Possible presence of stray dogs, any known canine leishmaniasis case in domestic/stray dogs around the region and the history of insecticide applications in the region (if known) were parameters taken into consideration.

**Figure 1:**
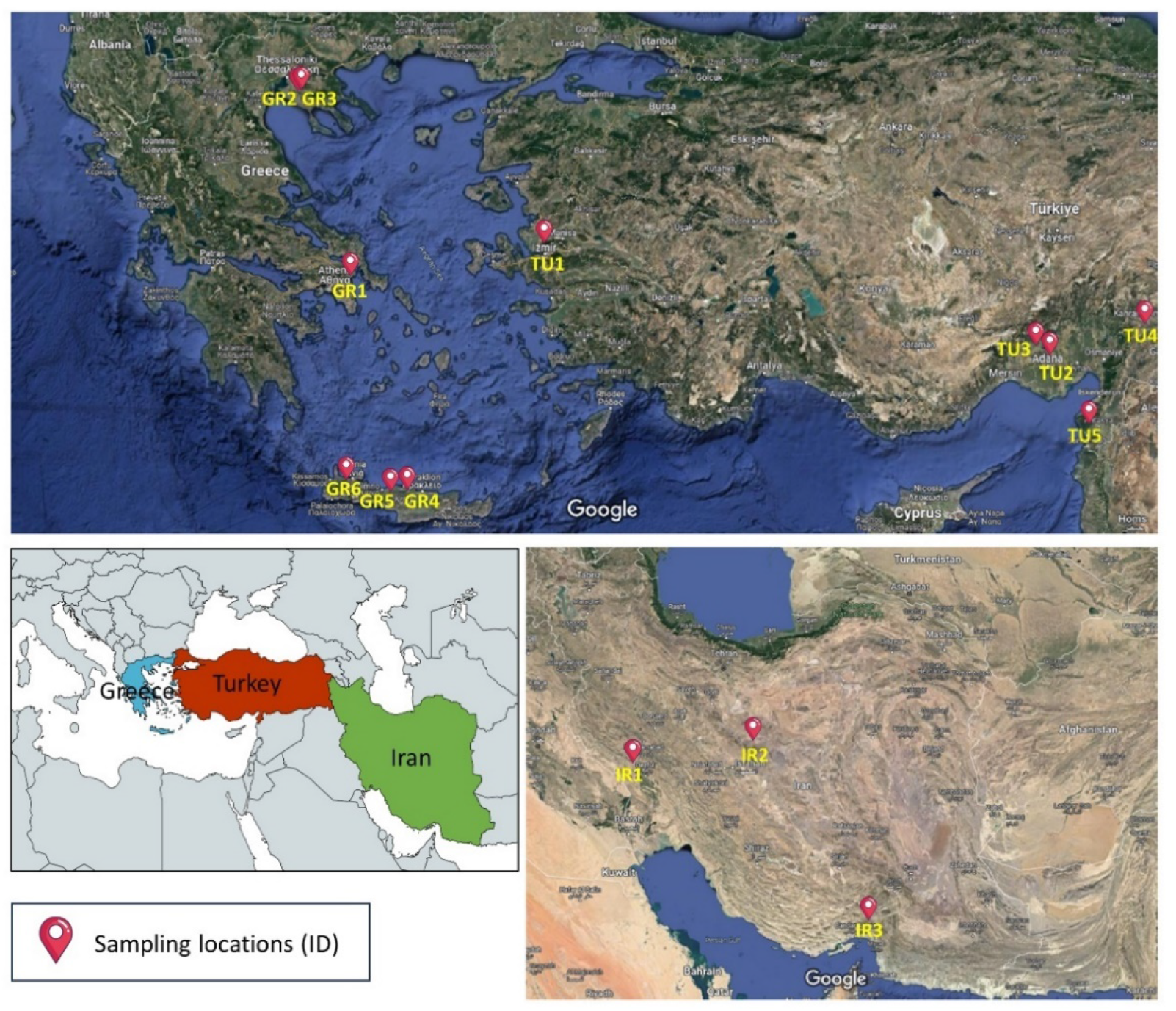
Sand fly sampling locations in Greece (GR1-4), Turkey (TU1-5) and Iran (IR1-3). Surveyed countries’ screenshots were obtained from Google Maps (accessed on 29 April, 2024), and the base layer of the lower left panel was created in mapchart.net.

**Table 1:**
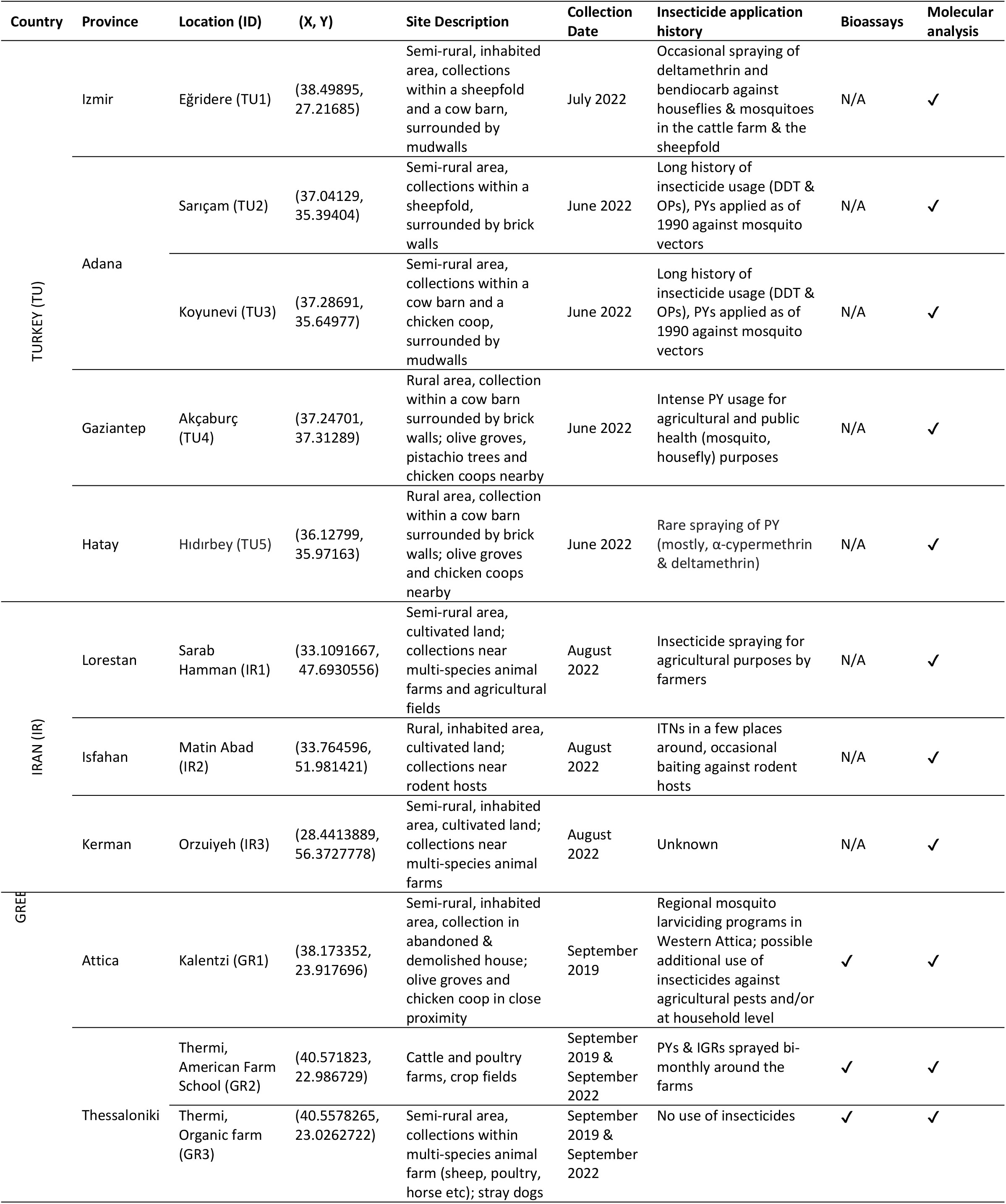
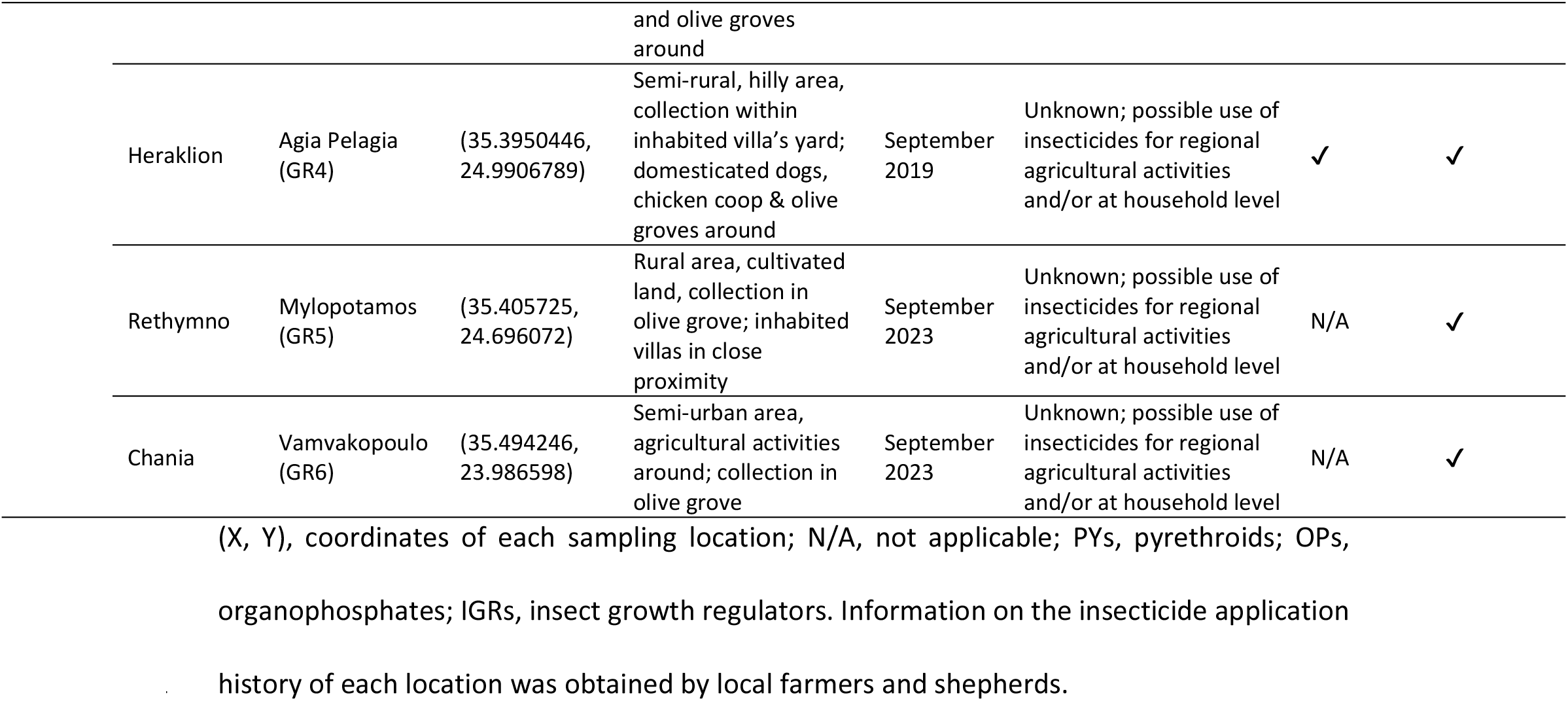
Description of sand fly sampling locations in Greece, Turkey and Iran, their insecticide application history (if known), and the type of analyses (phenotypic and/or molecular) each collected population was subjected to.

| Country | Province | Location (ID) | (X, Y) | Site Description | Collection Date | Insecticide application history | Bioassays | Molecular analysis |
| --- | --- | --- | --- | --- | --- | --- | --- | --- |
| TURKEY (TU) | Izmir | Eğirdere (TU1) | (38.49895, 27.21685) | Semi-rural, inhabited area, collections within a sheepfold and a cow barn, surrounded by mudwalls | July 2022 | Occasional spraying of deltamethrin and bendiocarb against houseflies & mosquitoes in the cattle farm & the sheepfold | N/A | ✓ |
|  | Adana | Sarıçam (TU2) | (37.04129, 35.39404) | Semi-rural area, collections within a sheepfold, surrounded by brick walls | June 2022 | Long history of insecticide usage (DDT & OPs), PYs applied as of 1990 against mosquito vectors | N/A | ✓ |
|  |  | Koyunevi (TU3) | (37.28691, 35.64977) | Semi-rural area, collections within a cow barn and a chicken coop, surrounded by mudwalls | June 2022 | Long history of insecticide usage (DDT & OPs), PYs applied as of 1990 against mosquito vectors | N/A | ✓ |
|  | Gaziantep | Akçaburç (TU4) | (37.24701, 37.31289) | Rural area, collection within a cow barn surrounded by brick walls; olive groves, pistachio trees and chicken coops nearby | June 2022 | Intense PY usage for agricultural and public health (mosquito, housefly) purposes | N/A | ✓ |
| | Hatay | Hıdırbey (TU5) | (36.12799, 35.97163) | Rural area, collection within a cow barn surrounded by brick walls; olive groves and chicken coops nearby | June 2022 | Rare spraying of PY (mostly, α-cypermethrin & deltamethrin) | N/A | ✓ |
| IRAN (IR) | Lorestan | Sarab Hamman (IR1) | (33.1091667, 47.6930556) | Semi-rural area, cultivated land; collections near multi-species animal farms and agricultural fields | August 2022 | Insecticide spraying for agricultural purposes by farmers | N/A | ✓ |
|  | Isfahan | Matin Abad (IR2) | (33.764596, 51.981421) | Rural, inhabited area, cultivated land; collections near rodent hosts | August 2022 | ITNs in a few places around, occasional baiting against rodent hosts | N/A | ✓ |
|  | Kerman | Orzuiyeh (IR3) | (28.4413889, 56.3727778) | Semi-rural, inhabited area, cultivated land; collections near multi-species animal farms | August 2022 | Unknown | N/A | ✓ |
| GREE | Attica | Kalentzi (GR1) | (38.173352, 23.917696) | Semi-rural, inhabited area, collection in abandoned & demolished house; olive groves and chicken coop in close proximity | September 2019 | Regional mosquito larviciding programs in Western Attica; possible additional use of insecticides against agricultural pests and/or at household level | ✓ | ✓ |
|  | Thessaloniki | Thermi, American Farm School (GR2) | (40.571823, 22.986729) | Cattle and poultry farms, crop fields | September 2019 & September 2022 | PYs & IGRs sprayed bi-monthly around the farms | ✓ | ✓ |
|  |  | Thermi, Organic farm (GR3) | (40.5578265, 23.0262722) | Semi-rural area, collections within multi-species animal farm (sheep, poultry, horse etc); stray dogs | September 2019 & September 2022 | No use of insecticides | ✓ | ✓ |

|  |  |  |  |  |  |  |  |
| --- | --- | --- | --- | --- | --- | --- | --- |
|  |  |  | and olive groves around |  |  |  |  |
| Heraklion | Agia Pelagia (GR4) | (35.3950446, 24.9906789) | Semi-rural, hilly area, collection within inhabited villa's yard; domesticated dogs, chicken coop & olive groves around | September 2019 | Unknown; possible use of insecticides for regional agricultural activities and/or at household level | ✓ | ✓ |
| Rethymno | Mylopotamos (GR5) | (35.405725, 24.696072) | Rural area, cultivated land, collection in olive grove; inhabited villas in close proximity | September 2023 | Unknown; possible use of insecticides for regional agricultural activities and/or at household level | N/A | ✓ |
| Chania | Vamvakopoulo (GR6) | (35.494246, 23.986598) | Semi-urban area, agricultural activities around; collection in olive grove | September 2023 | Unknown; possible use of insecticides for regional agricultural activities and/or at household level | N/A | ✓ |

(X, Y), coordinates of each sampling location; N/A, not applicable; PYs, pyrethroids; OPs, organophosphates; IGRs, insect growth regulators. Information on the insecticide application history of each location was obtained by local farmers and shepherds.

In Greece, sand flies were collected overnight using CDC light traps baited with dry ice, set before 5 pm and removed at 7 am the day after (for two to ten consecutive nights). The collection bags were transferred from the field to the laboratory. In Iran, a hand aspirator was used to collect sand flies at different time intervals between 8 pm and 5 am, and, afterwards, the samples were transferred from the field to the laboratory using cages. Both collection methods, i.e. CDC light traps and mouth aspirator, were combined in Turkish regions.

After field collections from Greece (in Kalentzi GR1, Thermi GR2 and GR3, and Agia Pelagia GR4), we proceeded to CDC bioassays, excluding male and engorged female sand flies (that were only included in the following molecular analyses). Upon bioassay completion, females were stored in absolute ethanol for subsequent molecular analyses. For the remaining sampling locations in the country (Vamvakopoulo GR5 and Mylopotamos GR6) and in all locations in Iran and Turkey, all the collected specimens were stored in ethanol for molecular analyses.

### 2. Genomic DNA extraction from sand flies

Genomic DNA (gDNA) was extracted from individual sand flies, using the DNazol reagent, according to the manufacturer’s instructions (Invitrogen, Carlsbad, CA, USA). The DNA quantity was assessed by NanoDrop 2000c spectrophotometer.

### 3. Molecular identification of sand fly species

Discrimination of sand fly species relied on PCR amplification of a 700 bp mitochondrial cytochrome oxidase subunit I (*COI*) genomic fragment, using primers LCO1490 and HCO2198 [32] and *Taq* DNA polymerase (EnzyQuest, Heraklion, Greece) on approximately 10-20 ng of gDNA template. The applied thermal protocol was as follows: 94°C for 2 min, 35 cycles x [94°C for 45 sec, 50°C for 30 sec, 72°C for 45 sec], 72°C for 10 min. After agarose gel visualisation of a small PCR product quantity (5 ul), the rest was purified using the Nucleospin PCR & Gel Clean- Up Kit (Macherey Nagel, Dueren, Germany) and, then, subjected to Sanger sequencing (GENEWIZ, Azenta Life Sciences, Germany) with the LCO1490 primer and BLASTn analysis.

### 4. Genotyping of mutations in the voltage-gated sodium channel (*vgsc*)

The presence of *kdr* mutations L1014F and L1014S, associated with resistance to pyrethroids and previously detected in sand flies and other insect species populations, was monitored in individual sand flies by genotyping the *vgsc* domain IIS6, containing codon 1014 (*Musca domestica* numbering). The genomic sequence was amplified by *Taq* DNA polymerase (EnzyQuest, Heraklion, Greece) using primers Vssc8F and Vssc1bR, as described in Gomes et al., 2017 [12], on approximately 10-20 ng of gDNA template. The reaction’s thermal conditions were: 94°C for 2 min, 35 cycles x [94°C for 45 sec, 56°C for 30 sec, 72°C for 30 sec], 72°C for 10 min. The approximately 400 bp generated PCR fragments, after visualization in agarose gel, were purified using the Nucleospin PCR & Gel Clean-Up Kit (Macherey Nagel, Dueren, Germany) and, then, subjected to Sanger sequencing (GENEWIZ, Azenta Life Sciences, Germany), using the Vssc8F primer. Sequences were analysed using the sequence alignment editor BioEdit 7.2.5 (https ://bioed it.softw are.infor mer.com/7.2/). Reference *vgsc* partial genomic sequences were obtained from GenBank for *Phlebotomus papatasi* (MH401419.1), *P. perfiliewi* (MG779187.1), *P. neglectus* (MG779192.1), *P. simici* (MG779189.1) and *P. tobbi* (MG779188.1).

### 5. Insecticide susceptibility bioassays

#### i. Standard CDC bottle bioassay

Female, non-engorged field-caught sand flies from Kalentzi (GR1), Thermi (GR2, GR3) and Agia Pelagia (GR4), Greece, were exposed to deltamethrin (pyrethroid insecticide), purchased as technical grade material (PESTANAL® analytical standard; Sigma-Aldrich, Darmstadt, Germany). Insecticide stock solutions were prepared in acetone and 250 ml Wheaton bottles were coated as described in the CDC guidelines [33]. The insecticide doses tested for the Greek populations were the CDC diagnostic for mosquito vectors; particularly, for deltamethrin 10 μg/ml (diagnostic dose for *Aedes* mosquitoes before 2021; applied to GR1- GR4 populations, in 2019 [20] and/or 0.75 μg/ml (current diagnostic dose for *Aedes* and *Culex*; applied to GR2 and GR3, in 2022). Two to six replicate bottles were tested for any given deltamethrin concentration (the number of replicates per population was relative to the number of collected sand flies) and a control bottle coated with acetone was always included. Alive and knocked-down sand flies in each bottle were recorded at time intervals of 5–15 minutes, for 1 hour exposure time. The percentage of mortality in each bottle was recorded after 24 hours of recovery; during that period, sand flies, transferred in plastic cups, were maintained in temperature and relative humidity (RH) conditions, as close to the optimal as possible (∼25°C, RH 70%), supplemented with 10% sucrose solution. The insecticide susceptibility status was determined by the mortality rate, according to CDC recommendations: 98–100% mortality indicates susceptibility; 80–97% suggests the possibility of resistance that requires further confirmation; and mortality < 80% denotes resistance. In cases where mortality (between 3– 5%) was recorded in the control bottle, mortality data were corrected using Abbott’s formula.

#### ii. Individual specimen glass vial bioassay (modified CDC bottle bioassay)

The objective behind generating this novel individual specimen bioassay protocol for sand flies is to detect any possible hint of tolerance/resistance to insecticides in single species, that might have been “diluted” during mixed species exposure. The protocol generated is a modified version of the CDC bottle bioassay protocol, adjusted in 9 ml round bottom, plain- end glass tubes of dimensions 12 x 100 mm (Fisherbrand^TM^). 24-hour viability of female sand flies inside these vials was tested using *Phlebotomus papatasi* laboratory colony (EBCL, USDA) specimens.

To apply insecticide doses equivalent to the CDC diagnostic ones (i.e. 1 ml of deltamethrin solution 0.75 μg/ml to coat a 250 ml Wheaton bottle), we needed 36 μl of 0.75 μg/ml deltamethrin solution for 9 ml glass vials’ coating, a volume which, during initial coating trials, was proved to be insufficient to fully coat the inner walls of them. Thus, we doubled the volume to 72 μl and prepared a 0.375 μg/ml deltamethrin solution. Similar modifications of the standard CDC bioassay protocol were previously described in Denlinger et al. 2015 [17] and Li et al. 2015 [18].

Control vials were coated with 72 μl of acetone and all vials were left to dry overnight in a shady place. The next day, using a mouth aspirator, individual females were transferred inside insecticide-treated and control vials, sealed with cotton. Each specimen’s survival was observed every 5 minutes until knock down and the knock-down time (KDT) was recorded. Afterwards, sand flies were stored individually in absolute ethanol for subsequent molecular identification of species.

## Results

### 1. Molecular identification of sand fly species

Six different *Phlebotomus* and two *Sergentomyia* species were recorded in the five regions of Greece (Table 2). In Attica, *P. neglectus* and *P. simici* were the most prevalent species (around 31% each), followed by *P. tobbi*. *Phlebotomus perfiliewi* was dominant in Thessaloniki, where exclusively detected, with a frequency over 40%, while more than 50% of the samples there were identified as *P. tobbi* or *P. simici*. In the Prefecture of Crete (Heraklion, Rethymno and Chania regions), *P. neglectus*, *P. simici* and *P. similis* denoted the highest occurrence among *Phlebotomus* species. *P. papatasi* was detected only in Thessaloniki and Rethymno, in frequencies below 5%. Lastly, *S. minuta* was present in all sampling regions of the country, peaking at Chania (37.5%), and *S. dentata* only in Attica and Thessaloniki, holding frequencies of 15.4% and 0.5% respectively.

**Table 2:**
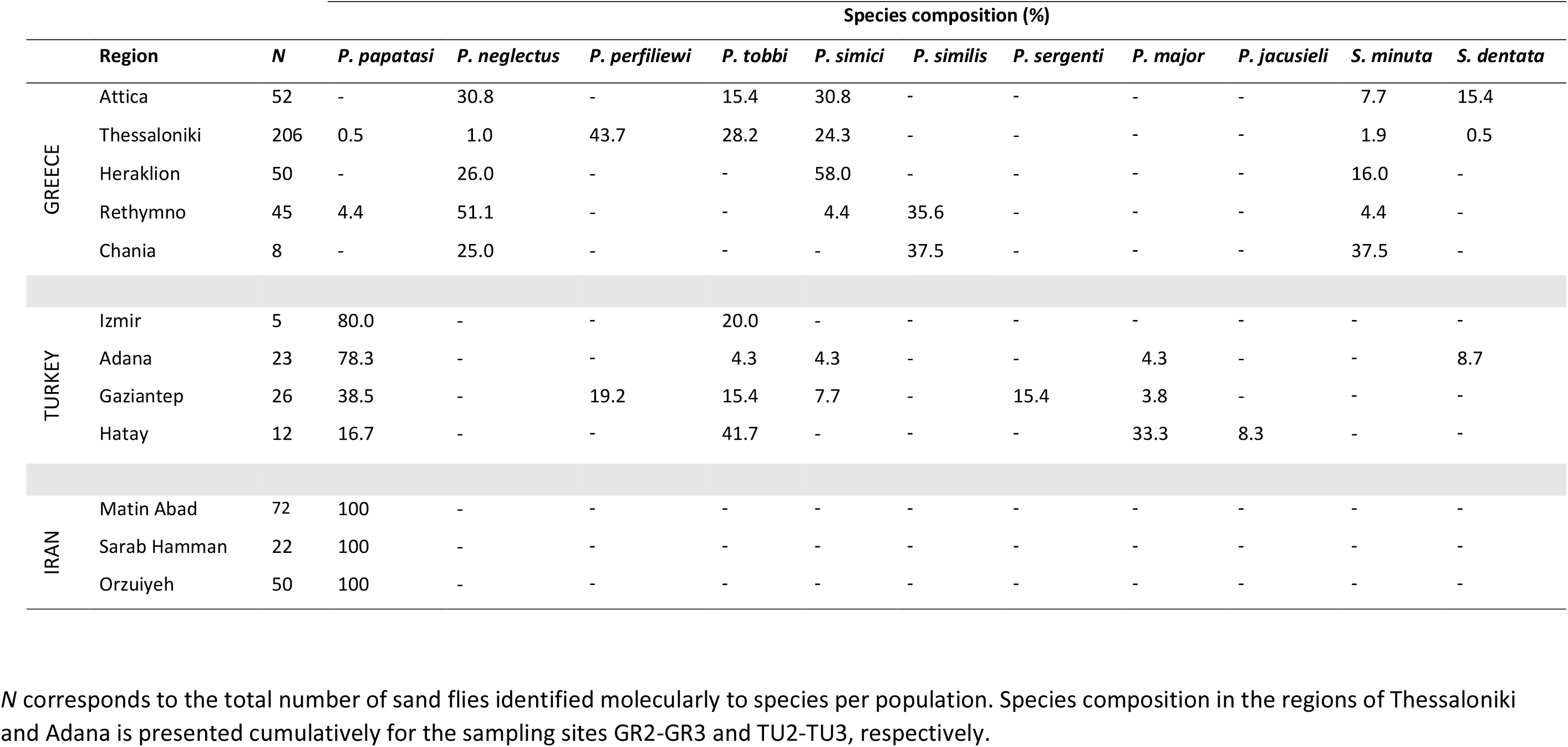
Sand fly species composition per sampling region.

*Phlebotomus* genus in Turkish populations was principally represented by *P. papatasi* (38.5 - 80%) in three out of four sampling locations. *Phlebotomus tobbi* frequencies ranged from 4.3 to 20% in most locations, except Hatay where they reached ∼40%. Other *Phlebotomus* species, particularly *P. perfiliewi*, *P. simici*, *P. sergenti* and *P. jacusieli*, were also present in some locations with frequencies below 20%. *P. major* was reported in all regions apart from Izmir (overall very low number of collected specimens in this region), reaching a maximum frequency of almost 35% in Hatay. *S. dentata* was the sole *Sergentomyia* species detected in Turkey (<10% in Adana) (Table 2).

In Iran, all collected samples were exclusively identified as *P. papatasi*.

### 2. Monitoring of pyrethroid resistance mutations *kdr* L1014F/S

*Phlebotomus* and *Sergentomyia* female and male samples from all regions were analyzed for the presence of *vgsc* mutations L1014F and L1014S, associated with pyrethroid resistance (Table 3).

**Table 3:**
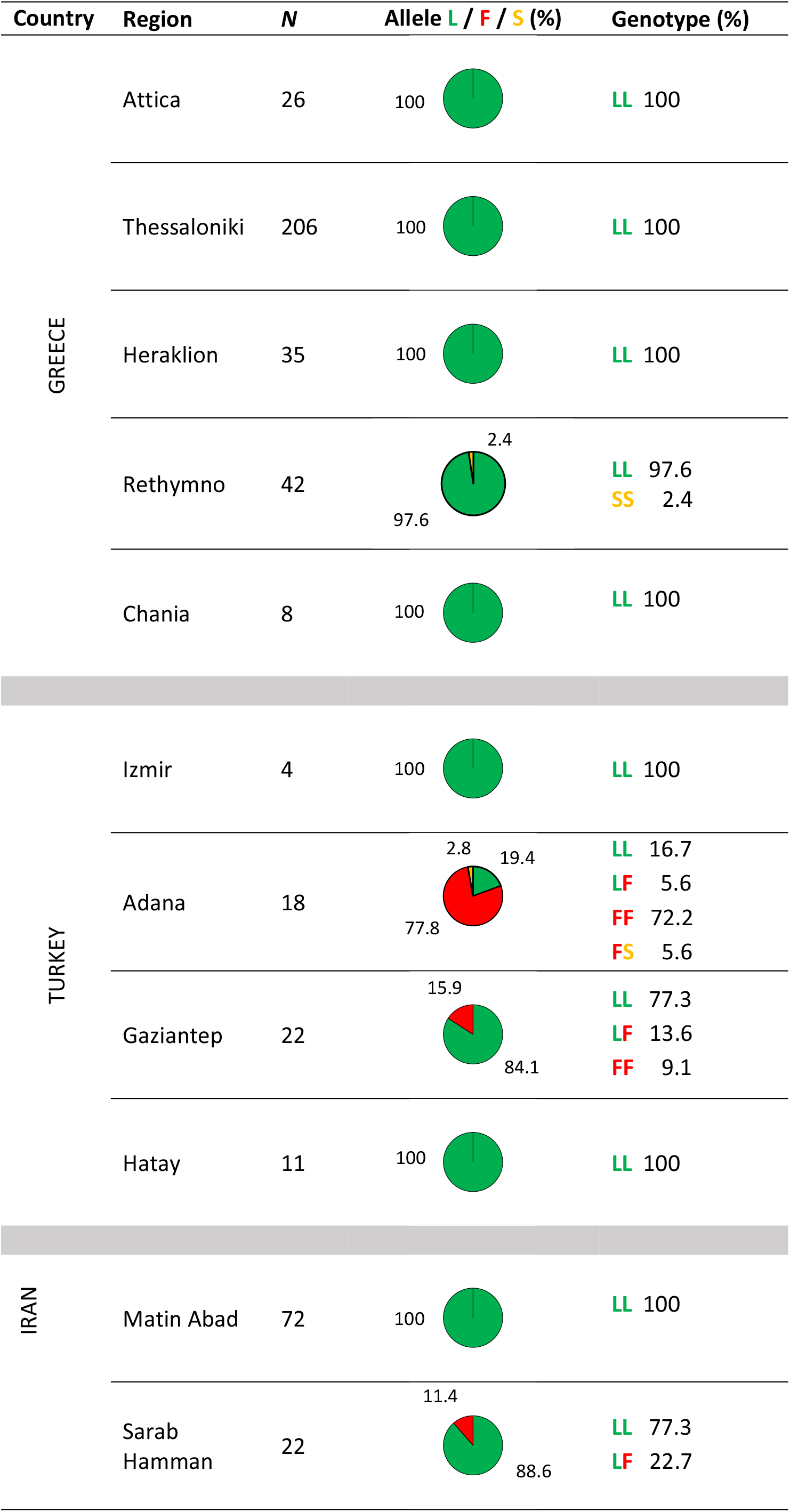
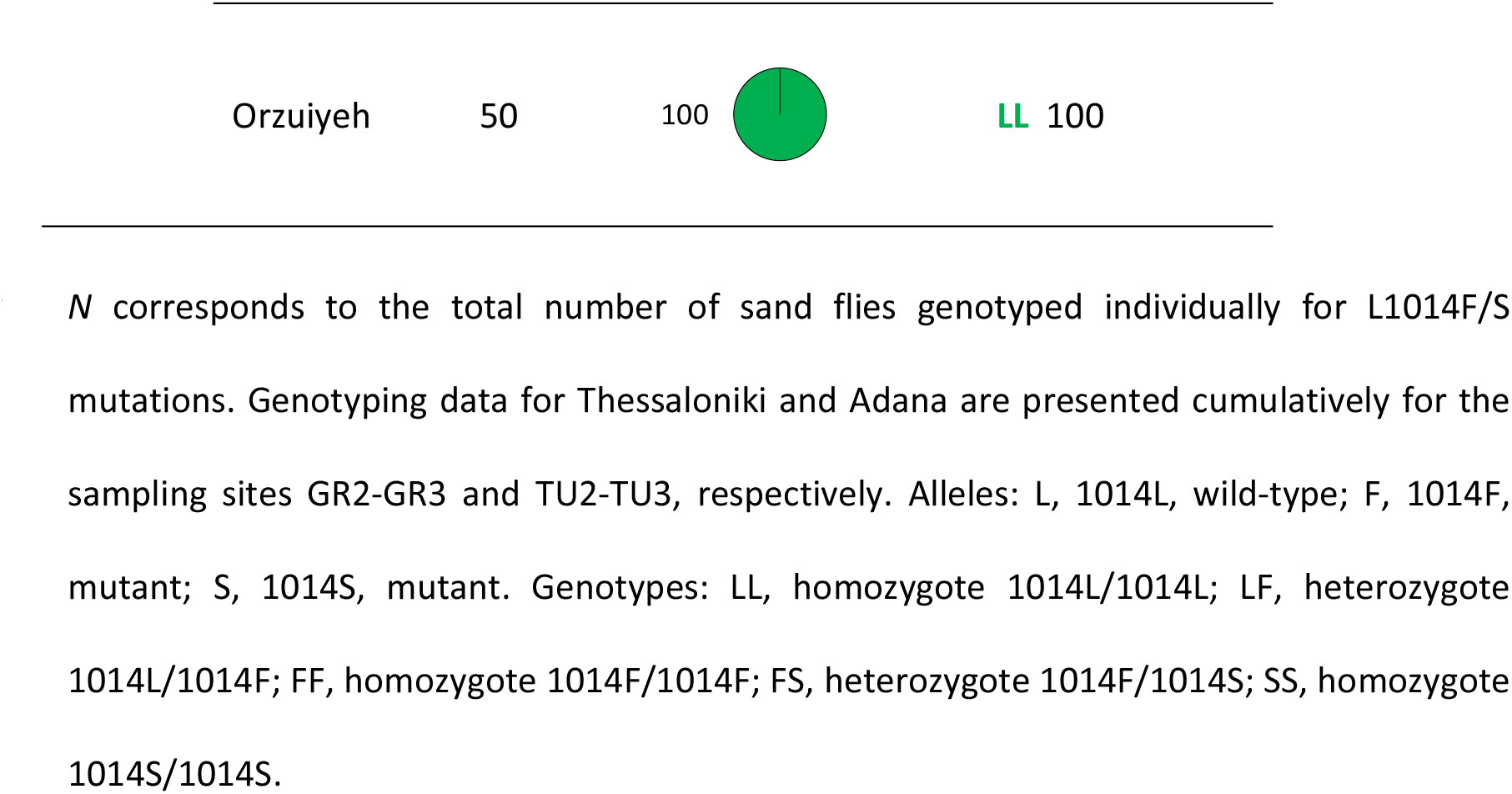
Molecular monitoring of *kdr* mutations L1014F/S in the collected sand fly populations.

Mutant allele 1014F (codon TTT) held a surprisingly high frequency, reaching almost 80% in Adana (locations TU2, TU3), Turkey, while occurring in homozygosity in 72.2% of the samples; near all 1014F/F homozygotes belonged to *P. papatasi*, except for one *S. dentata*. A single *P. tobbi* specimen harboured 1014F in heterozygosity. Mutation 1014S (serine encoded by TCA) was additionally recorded in Adana (frequency <3%), in a heterozygote (1014F/S) *P. papatasi* individual. Moreover, in Gaziantep (TU4), 1014F was the only mutant allele present, in a frequency of 15.9%, with two *P. papatasi* homozygotes and three 1014L/F heterozygote samples (two *P. papatasi* and one *P. perfiliewi)*. All samples from Izmir (TU1) and Hatay (TU5) were wild-type (1014L/1014L; TTA for leucine).

In Sarab-e Hamman (IR1), the only *kdr*-positive population from Iran, 5 out of the 22 genotyped *P. papatasi* individuals carried 1014F allele in heterozygosity (genotype frequency 11.4%).

In Greece, no population harboured mutation L1014F, while only one *P. papatasi* specimen from Rethymno (GR5) was homozygote for L1014S (frequency 2.4%).

### 3. Insecticide susceptibility bioassays

#### i. Standard CDC bottle bioassays

Mixed species populations from Greece, i.e. Kalentzi (GR1, Attica), Thermi (GR2 and GR3, Thessaloniki) and Agia Pelagia (GR4, Heraklion), collected in 2020, were exposed to deltamethrin 10 μg/ml for 1 hour. Fifty percent of all populations were knocked down within the first 20 minutes of exposure, and no significantly different knock-down rate was noted among them (Figure 2A). Mortality recorded after 24 hours was 100% for all four populations, indicating susceptibility, according to CDC guidelines (Figure 2B).

**Figure 2.**
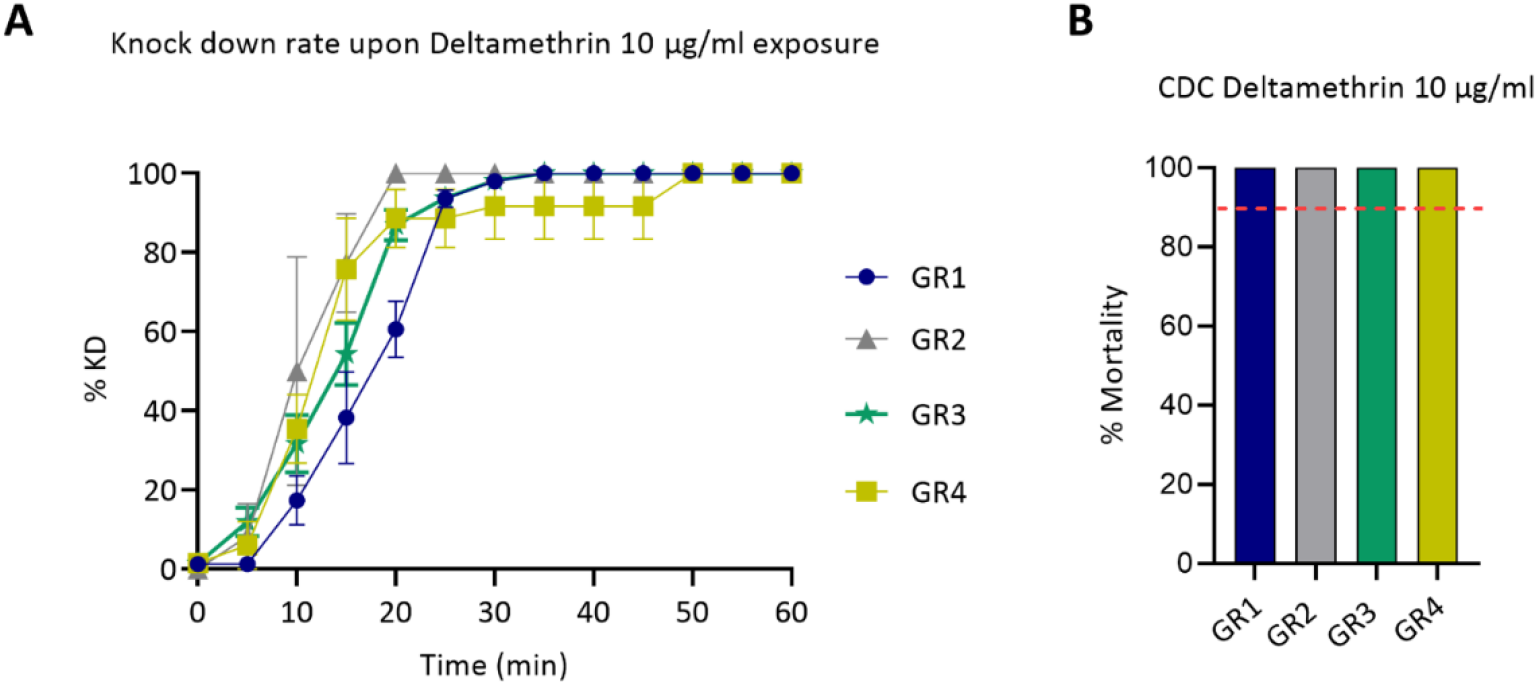
Response of GR1-GR4 mixed species populations from Greece, to deltamethrin 10 μg/ml in CDC bottle bioassays. A) Knock-down rate recorded during one hour exposure to deltamethrin. Results are expressed as the mean knock down percentage (%) and bars represent SEM (*n*=2 to 5 biological replicates). B) Percentage (%) of mortality recorded 24 hours post exposure. Results are expressed as the mean mortality percentage (%). The red dotted line corresponds to the 90% mortality cut off indicating resistance, according to CDC guidelines.

During collection year 2022, to exclude the possibility of any upcoming resistance/ tolerance in the regularly pyrethroid-treated GR2 population (American Farm School, Thermi), we exposed this population comparatively to its proximal GR3 (Organic Farm, Thermi), to a much lower deltamethrin dose, 0.75 μg/ml, for 1 hour. Both populations displayed an almost identical response and, as expected, a slower knock down rate compared to the 10 μg/ml exposure (Figure 3A), yet 100% mortality (Figure 3B).

**Figure 3.**
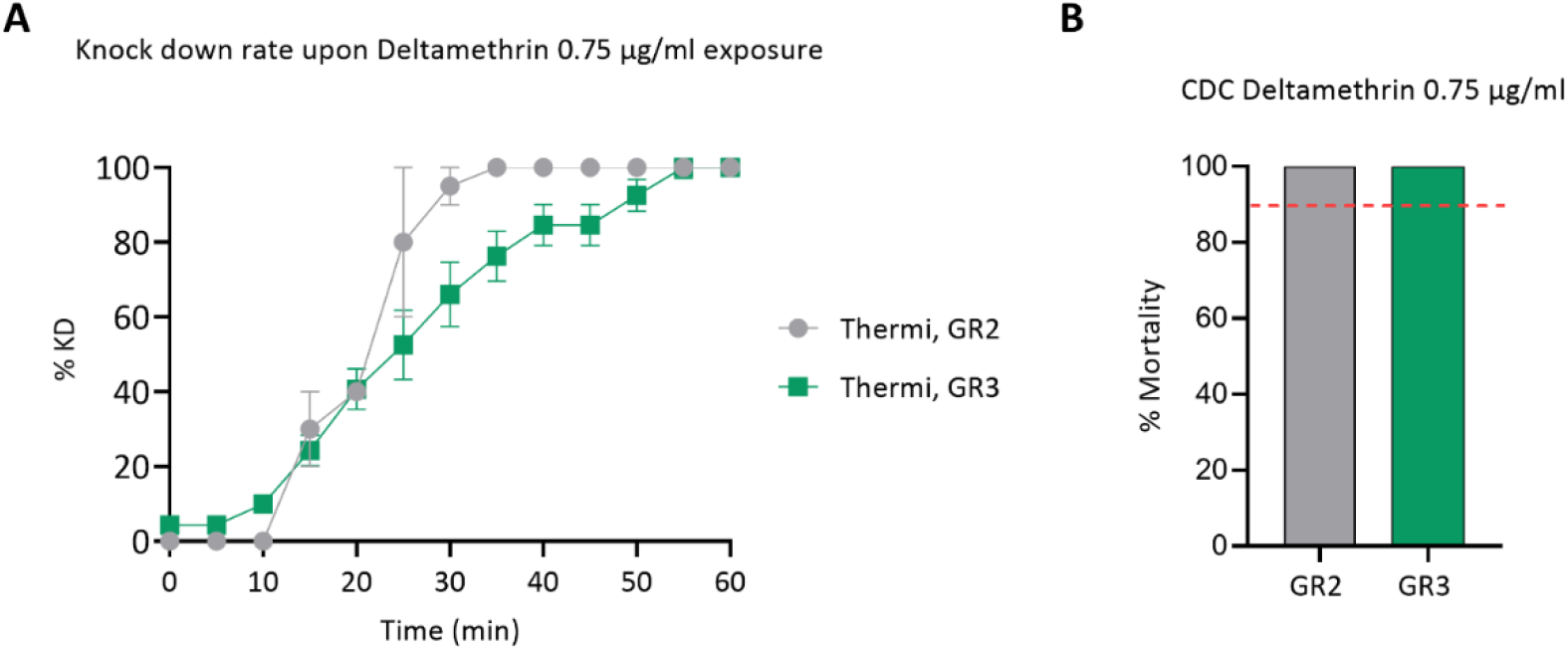
Response of GR2 and GR3 mixed species populations from Thermi, Thessaloniki, to deltamethrin 0.75 μg/ml in CDC bottle bioassays. A) Knock-down rate recorded during one hour exposure to deltamethrin. Results are expressed as the mean knock down percentage (%) and bars represent SEM (*n*=2 to 5 biological replicates). B) Percentage (%) of mortality recorded 24 hours post exposure. Results are expressed as the mean mortality percentage (%). The red dotted line corresponds to the 90% mortality cut off indicating resistance, according to CDC guidelines.

#### ii. Individual specimen glass vial bioassays

Specimens collected, during summer 2022, from locations GR2 and GR3, in Thermi- Thessaloniki (considered here as a single population, given their close proximity, their similar species composition and the absence of any significant difference in their knock-down rate, in CDC deltamethrin 0.75 and 10 μg/ml bioassays; see Fig.2 and Fig.3), were exposed individually to deltamethrin 0.75 μg/ml. The insecticide dose was chosen based on the previous observations during CDC bioassays, where sand flies from the same locations exhibited a “smoother” knock-down rate when exposed to 0.75 μg/ml, making any differential response more noticeable.

Post deltamethrin exposure and molecular identification of species, it was revealed that the mean KDT of *P. simici* samples was approximately 60 min, significantly higher from *P. perfiliewi* KDT 35.3 min (**, *p* = 0.001). *P. tobbi* denoted an intermediate mean KDT of 43.3 min, not significantly different from both *P. perfiliewi* and *P. simici* (Figure 4).

**Figure 4.**
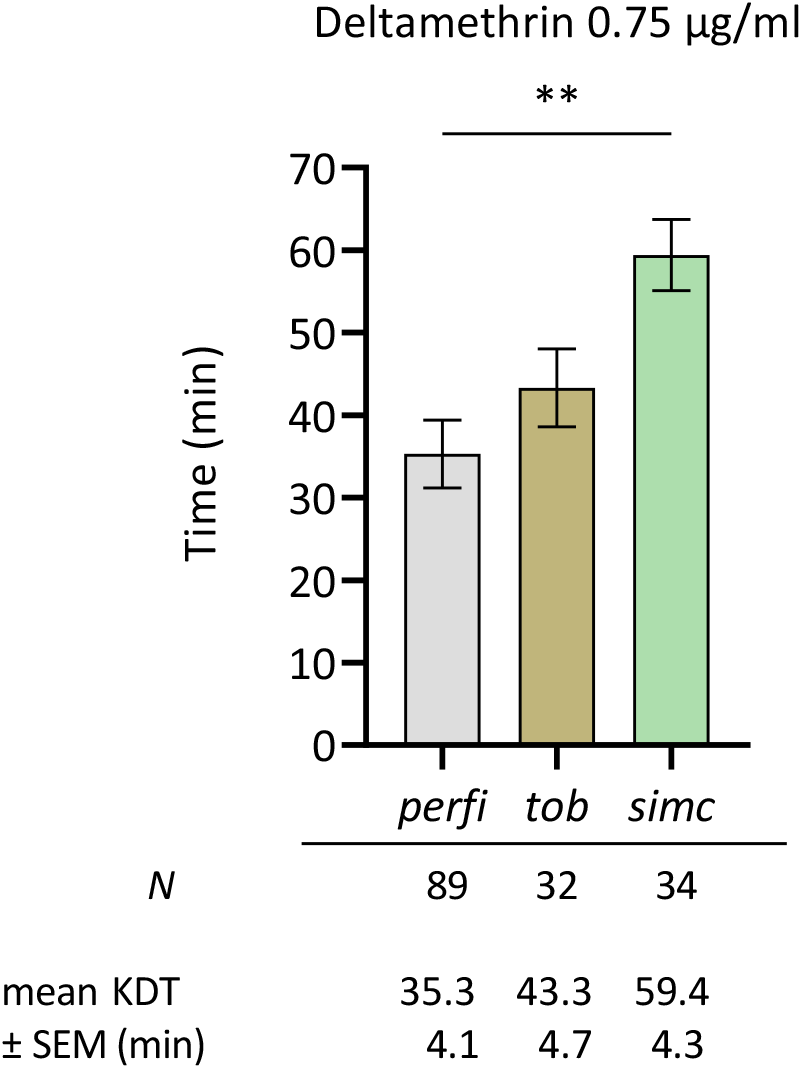
Mean knock down time (KDT) of three *Phlebotomus* species, i.e. *perfi, P. perfiliewi*; *tob, P. tobbi*; *simc*, *P. simici*, from Thermi, Greece, upon exposure to deltamethrin 0.75 μg/ml, in individual glass vial bioassays. *N* corresponds to the total of individually exposed female specimens per species, cumulatively from populations GR2 and GR3. Bars represent SEM. Results were statistically analysed using one-way ANOVA; **, *p*=0.001.

## Discussion

Leishmaniasis mitigation efforts, largely rely on sand fly control by means of insecticide-based tools, are threatened by the development of insecticide resistance (IR) phenotypes in vectors’ wild populations [34]. This study represents an attempt to address this critical knowledge gap in sand flies by developing and applying entomological and IR profiling tools, in wild populations from three leishmaniasis endemic countries of the Mediterranean basin and the Middle East with limited monitoring data, i.e. Greece, Turkey and Iran. Our on-focus sampling locations were in (semi-)rural regions, with agricultural and/or livestock farming, in the vast majority of which insecticides are applied mainly for crop protection and/or occasionally for public health purposes (e.g. targeting mosquito vectors), at regional and/or at farm/household level.

Molecular identification of species in the collected populations appeared largely in line with previously reported data [29, 35–39]. *Phlebotomus neglectus*, *P. tobbi*, and *P. simici* had the most extensive distribution in the mainland of Greece and in the Island of Crete (Heraklion, Rethymno and Chania), while *P. perfiliewi* and *P. similis* appeared exclusively in Thessaloniki and Crete, respectively, in accordance with data generated by Chaskopoulou et al 2016 [35] and Dvorak et al 2020 [36]. *Sergentomyia minuta* was present in all Greek locations, as expected, in moderate to low frequencies. In Turkey, *P. papatasi* was the most prevalent species in three out of four surveyed Provinces, followed by *P. tobbi* and *P. major,* conforming with Kasap et al 2019 [29]. *Phlebotomus perfiliewi* and *P. sergenti* were detected only in Gaziantep. In Iran, *Phlebotomus papatasi* was the sole species identified, representing the dominant cutaneous leishmaniasis vector across the whole country [31]. It needs to be clarified though that our data neither reflect the integrated species composition of each region nor the seasonal abundance of species, since we focused on specific habitats (as described in Table 1), where we conducted a few consecutive night samplings, rather than systematic entomological surveillance. Nonetheless, it is encouraging to observe that the sand fly species population structure resulting from these targeted and relatively limited collection interventions is in agreement with data reported from more intensive surveillance efforts.

Interestingly, pyrethroid resistance mutations were recorded in all three surveyed countries, albeit in different frequencies. In Adana and Gaziantep, regions with long history of pyrethroid usage in Turkey, L1014F was recorded mainly in *P. papatasi*, particularly prevalent in the former region (frequency >70%). This mutation was previously identified in same species samples from Sanliurfa [40], an area close to our sampling regions in the south-eastern part of the country. In addition to that, mutation L1014S was noted for the first time in the country, in one 1014S/1014F *P. papatasi* sample from Adana, while in Izmir and Hatay regions, no mutation was recorded, yet the sample sizes were very small. *Kdr* L1014S and L1014F occurred likewise in *P. papatasi* specimens from Rethymno, Greece, and Sarab Hamman, Iran, respectively, albeit at frequencies below 12%. To the best of our knowledge, this is the first study to detect *kdr* mutations in wild sand fly populations from Greece and Iran, although certain regions of each country have been surveyed before [40–42].

The occurrence of resistance mutations, here, has probably followed the pyrethroid and/or DDT usage (as *kdr* mutations confer cross-resistance) implemented in the respective rural locations (e.g. regions Adana, Gaziantep, Sarab Hamman, etc), for agricultural protection and/or mosquito control (relevant information on past and/or current insecticide applications in each sampling location was obtained by local farmers and shepherds). Molecular markers could have a significant predictive value and ease the early detection of developing resistance [43]. As of 2017, several studies have monitored *kdr* mutations in sand fly populations from eight leishmaniasis endemic countries, in the Mediterranean basin, the Middle East and south- eastern Asia [11, 44–46]. In all of the surveyed countries, apart from Italy [47], i.e. India, Sri Lanka, Bangladesh, Iran, Armenia, Turkey and Greece, L1014F and/or L1014S were indeed present. Such findings highlight that systematic regional IR surveillance is necessary, as pyrethroid resistance might be spreading in sand fly populations.

Besides molecular diagnostics, CDC bottle bioassays were deployed to assess the susceptibility to deltamethrin of mixed sand fly species populations collected from Kalentzi- Attica (GR1), Thermi- Thessaloniki (GR2, GR3), and Agia Pelagia- Heraklion (GR4), Greece. Extrapolating the diagnostic doses of mosquitoes (10 μg/ml and, later, 0.75 μg/ml), they displayed a “susceptible” phenotype, with no notable difference in their knock-down rate during exposure, concurring with the absence of associated *kdr* mutations. Even at the lowest tested dose, the American Farm School and the nearby Organic Farm populations, sharing the same species structure, yet polar opposite insecticide exposure records, responded in a same manner, possibly implying that no apparent resistance to deltamethrin has been developed over the years in the regularly pyrethroid-sprayed AFS population. Nonetheless, we need to consider, that sand fly bioassay results using mosquito discriminating doses might be difficult to interpret [11, 16] and might have limited operational value for vector control, as any upcoming resistance is likely to pass unnoticed in case the tested dose is too high for sand flies *per se*.

Another critical point in phenotypic monitoring of IR is that exposing a mixed species population in a single CDC bottle (or WHO tube) might conceal possible divergent phenotypes of single species [48]. Proceeding to individual oviposition of field-caught females to create laboratory-reared isofemale progenies and, then, to bioassays, entails certain difficulties and risks, as discussed by Shirani-Bidabadi et al., 2020 [19]. Herein, to identify any obscured signs of tolerance or resistance to insecticides in single species, an alternative bioassay protocol to expose single sand fly specimens in insecticide-coated glass vials right after field collections is proposed. Applying this protocol to the populations GR2 and GR3 collected from Thermi, Greece, revealed that *Phlebotomus* species might indeed hold differing responses to insecticides. Particularly, *P. simici* exhibited an almost 2x-fold higher KDT than *P. perfiliewi* upon treatment with 0.75 μg/ml deltamethrin (mean KDT 59.4 vs 35.3 min). *P. simici*, a suspected *L. infantum* vector and among the most dominant species in our collections from the mainland of Greece, is well-established in the eastern Mediterranean basin and was recently recorded for the first time in Austria, raising concerns on its possible northward geographical expansion [49]. Even though we perceive that this is a time-consuming and technically difficult methodology, especially when in field conditions, it can potentially provide valuable information for effective sand fly control in relevant eco-epidemiological settings.

Nevertheless, it is essential to point out that resistance traits can be focally distributed, following the insecticide selection pressure regimes in each location, and thus probably rare to detect, especially given the mixed species composition [50], and the confined spatial movements that certain sand fly species might display [51]. Hence, we cannot rule out the possibility that target-site resistance traits (or detoxification-based, that were not examined here) may similarly occur in the rest of the analysed collections, especially those represented by a small sample size (such as GR2 and GR6 in Greece, and TU1 and TU5 in Turkey).

Monitoring the development of IR in sand flies is a core-element of integrated vector management (IVM), providing evidence-based guidance to vector control campaigns in an eco-epidemiologically context-specific manner. Following the paradigm of malaria mosquito control, tailoring *Leishmania* infection surveillance systems to IVM would stand as a proactive approach to minimize leishmaniasis incidence [34].

## Conclusion

In short, this study significantly enriches the IR status map of sand fly populations in regions of the Mediterranean basin and the Middle East, poorly monitored thus far. Molecular analyses of pyrethroid resistance in populations collected from Greece, Turkey and Iran, focused on *kdr* mutations, lately often reported in leishmaniasis endemic regions. The occurrence of L1014F/S mutations is more prevalent in Turkey and in lower frequencies, yet recorded for the first time, in Greece and Iran, implying emerging resistance. Phenotypic assessment of some mixed species populations from Greece against deltamethrin indicated susceptibility, extrapolating though the diagnostic doses for mosquitoes, an approach that usually leads to difficult-to-evaluate/-interpret results. Added to these, by establishing a novel bioassay protocol for individual specimens, we managed to show that sand fly species could display significantly different response to insecticides, highlighting the importance of resistance monitoring and regional vector control to be context- and species- specific. Given the limited arsenal of insecticide compounds appropriate for public health protection, sand fly control should be evidence-based, relying on regular species composition and IR monitoring, the latter encompassing both molecular and bioassay data, derived from sand fly- standardized protocols. Last but not least, there is an urgent need to generate relevant genomic resources for sand flies, that would facilitate the functional validation of new IR mechanisms, as well as the development of corresponding markers to diagnose target-site- as well as detoxification-based IR.

## Acknowledgments

This work was financially supported by the Hellenic Foundation for Research and Innovation (HFRI) under the 3rd Call for HFRI PhD Fellowships awarded to S.B. (Fellowship Number: 11078), by Fondation Santé through a research grant awarded to K.M., and also by the European Union’s Horizon Europe research and innovation programme under grant agreement number 101046133 (ISIDORe). L.R. has been supported by a Marie Curie Post Doctoral Fellowship (HORIZON-MSCA-2023-PF-01-01, project: 101152599).

